## Supplementary Figures and tables for "Dual role of FOXG1 in regulating gliogenesis in the developing neocortex via the FGF signalling pathway"

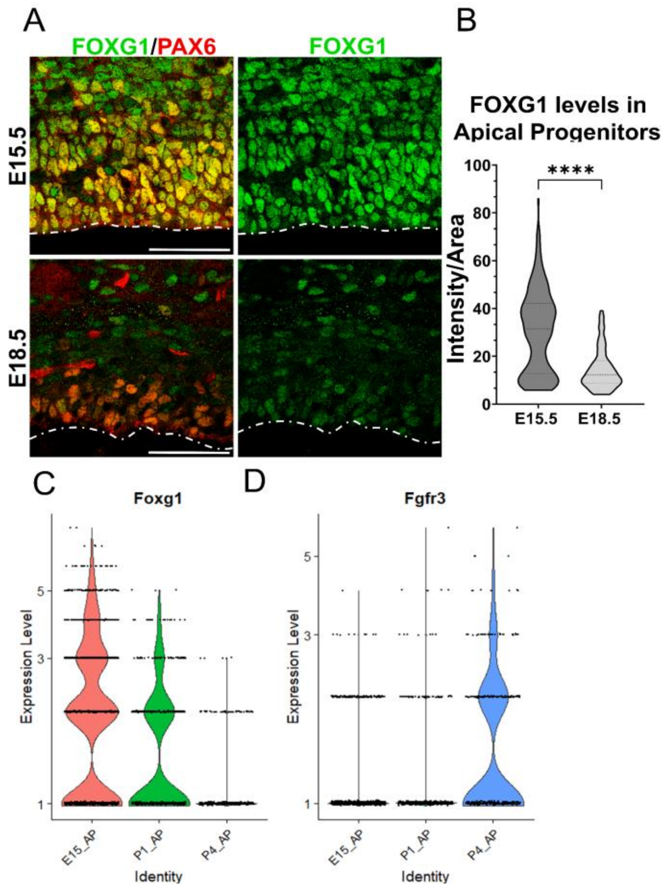


**Supplementary Figure S1: Complementary temporal regulation of *Foxg1* and *Fgfr3* in progenitors during the gliogenic period.**

(A) FOXG1 immunostaining at E15.5 and E18.5 in control brains together with apical progenitor marker PAX6. (B) Nuclear intensity quantification of FOXG1 in apical progenitors reveals a significant decrease in levels by E18.5, suggesting an endogenous mechanism for the transition of neurogenesis to gliogenesis in progenitors. n=350 cells from N=3 (biologically independent replicates). *Statistical test: Mann-Whitney Test. Scale bar: 50 µm.* (C, D) Violin plots depicting normalised gene expression levels from scRNA-seq data (76) for *Foxg1* (C) and *Fgfr3* (D) reveal complementary temporal dynamics for these genes in progenitors at E15.5, P1, and P4.

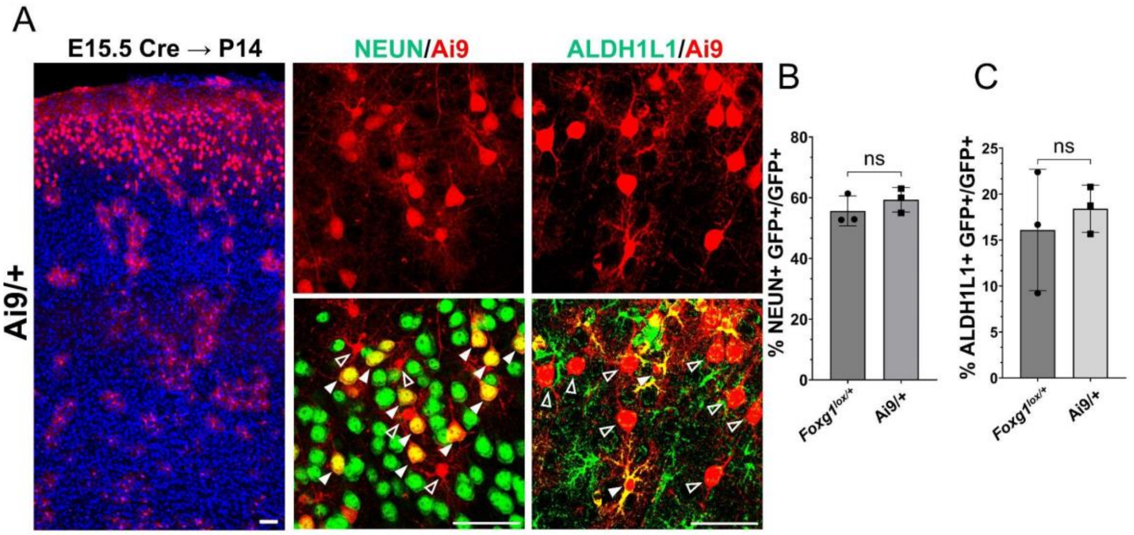


**Supplementary Figure S2: *Foxg1* haploinsufficiency at E15.5 does not lead to premature gliogenesis.**

This experiment aimed to examine whether there is a difference in baseline neurogenesis in *Foxg1*$^{+/+}$ versus *Foxg1*$^{lox/+}$ brains since the latter are used as controls in Figures 1 and 6. (A) Cre electroporation at E15.5 in control embryos carrying an *Ai9* reporter. Brains were harvested at P14. The leftmost panel in A is a low-magnification image, as the scale bar indicates. (B) Quantification of Ai9+ cells reveals similar proportions of these cells colocalise with NeuN in +/+ controls (72%) compared to *Foxg1*$^{lox/+}$*; GFP* $^{FRT}$ brains (67%). (C) Quantification of glial marker ALDH1L1 reveals similar proportions of Ai9+ cells colocalise with ALDH1L1 in +/+ brains (18%) compared to Foxg1$^{lox/+}$; GFP $^{FRT}$ brains (16%). No significant difference is observed between the two conditions. n=2790 cells from N=3 (biologically independent replicates) were examined over three independent experiments. *Statistical test: Unpaired t-test.*

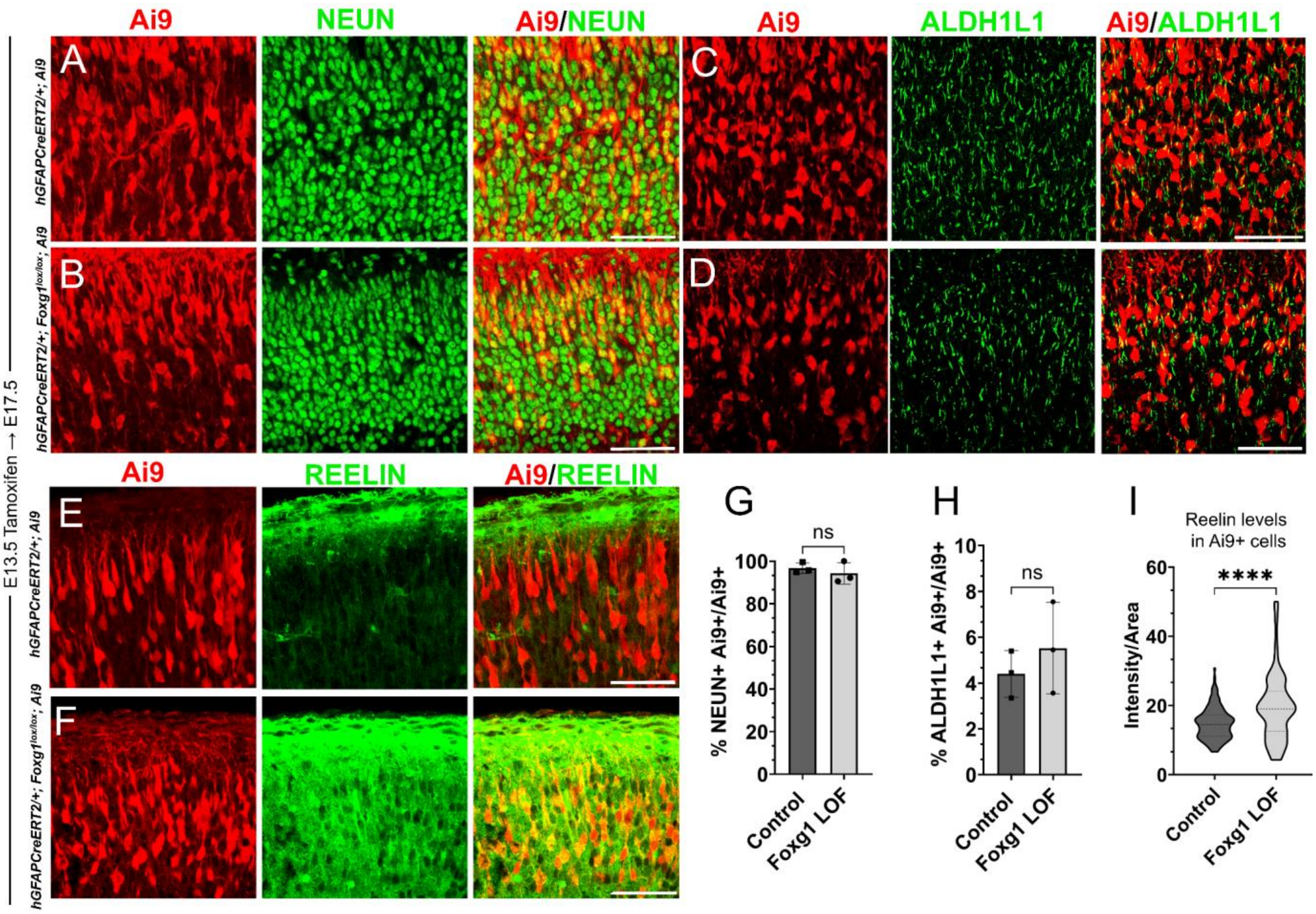


**Supplementary Figure S3: Loss of *Foxg1* at E13.5 does not lead to premature gliogenesis but results in increased REELIN.**

(A–F) *hGFAPCreERT2/+; Ai9* (Control) and *hGFAPCreERT2/+; Foxg1*$^{lox/lox}$*; Ai9* (*Foxg1* LOF) brains after tamoxifen administration at E13.5, followed by analysis at E17.5. (A, B, G) 96% of Ai9+ cells colocalised with NEUN in control brains and 95% in *Foxg1* LOF brains. (C, D, H) 4% of Ai9+ cells colocalised with ALDH1L1 in control brains and 5% in *Foxg1* LOF brains. (E, F, I) The average REELIN intensity per Ai9+ cell increased from 17% in control brains to 22% in *Foxg1* LOF brains. n= 2508 (Control), 2798 (*Foxg1* LOF) cells from N=3 brains (biologically independent replicates).

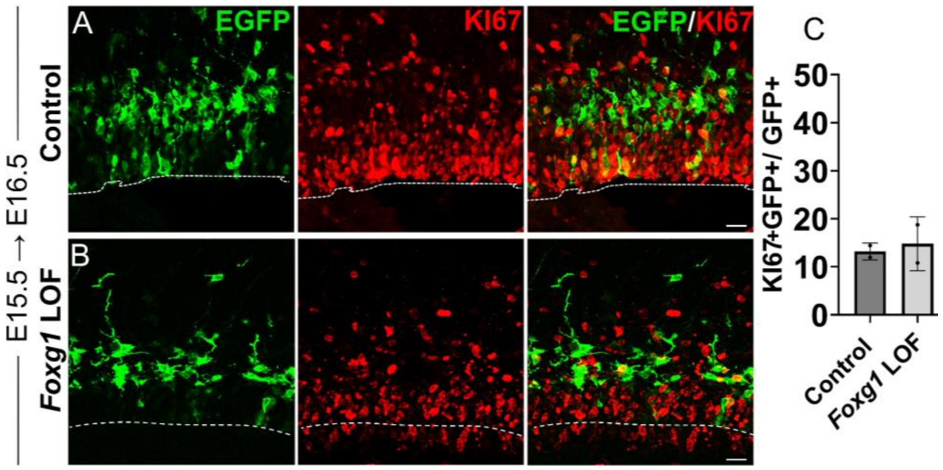


**Supplementary Figure S4: Loss of *Foxg1* from E15.5 does not lead to enhanced proliferation at E16.5**

Analysis of the proliferation marker KI67 in Control and *Foxg1* LOF E16.5 progenitors, one-day post-electroporation at E15.5. KI67 colocalises with comparable numbers of GFP+ cells in both Control (A; Cre in *Foxg1*$^{lox/+}$) and *Foxg1* LOF (B; Cre in *Foxg1*$^{lox/lox}$) brains. Quantifications are presented in C. n=403 cells (Control) and 663 cells (*Foxg1* LOF) from N=2 biologically independent experiments.

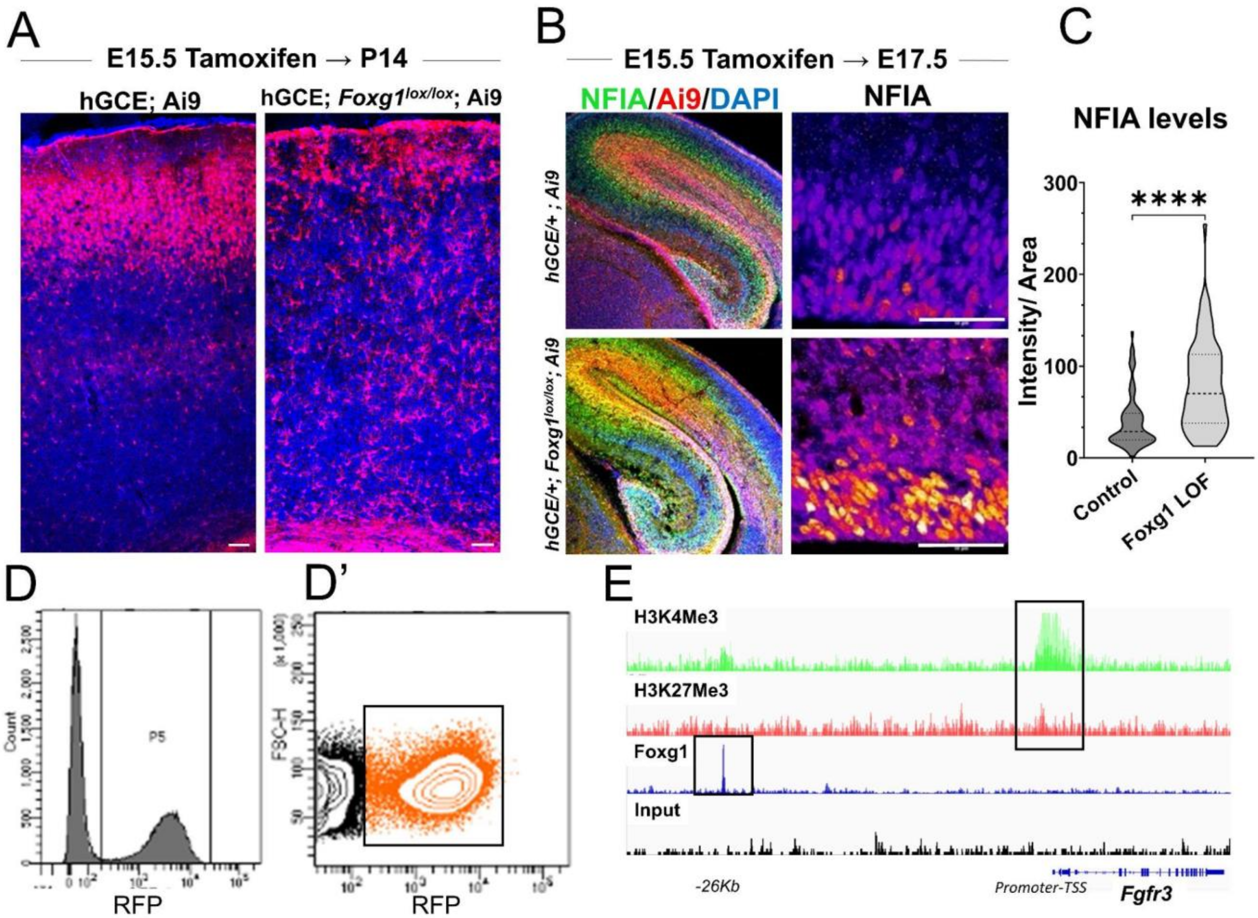


**Supplementary Figure S5: *hGFAPCreERT2; Ai9* (hGCE) line-based recapitulation of the premature gliogenesis phenotype and subsequent analysis.**

(A) *hGFAPCreERT2/+; Ai9* (hGCE*; Ai9)* and *hGFAPCreERT2/+; Foxg1* $^{lox/lox}$*; Ai9* (labeled hGCE *Foxg1*$^{lox/lox}$; *Ai9*) brains after tamoxifen administration at E15.5 and harvesting at at P14. The *Foxg1* LOF condition shows the absence of neurons and enhanced glia, consistent with the phenotype from the electroporation-mediated loss of *Foxg1*. This result is similar to that in Figure 1. (B, C) hGCE *Foxg1*$^{lox/lox}$; *Ai9* (tamoxifen at E15.5, harvested at E17.5). *Foxg1* LOF progenitors display a significant upregulation of gliogenic factor NFIA (n=310 (hGCE*; Ai9*) and 367 (hGCE *Foxg1*$^{lox/lox}$; *Ai9*) cells from N=3 biologically independent replicates). (D, D') Fluorescence-Activated Cell Sorting (FACS) plot of the Ai9+ cells obtained from control and hGCE *Foxg1*$^{lox/lox}$; *Ai9* brains (tamoxifen at E15.5, harvested at E17.5). Cells in the boxed cluster in D' were collected at E17.5 and purified for RNA sequencing. (E) Plot depicting the presence of both H3K4Me3 and H3K27Me3 (Bivalent) marks at the *Fgfr3* promoter region and the presence of a FOXG1 binding site at 26Kb upstream region of the gene. *Statistical test: Mann-Whitney Test. Scale bar: 50 µm.*

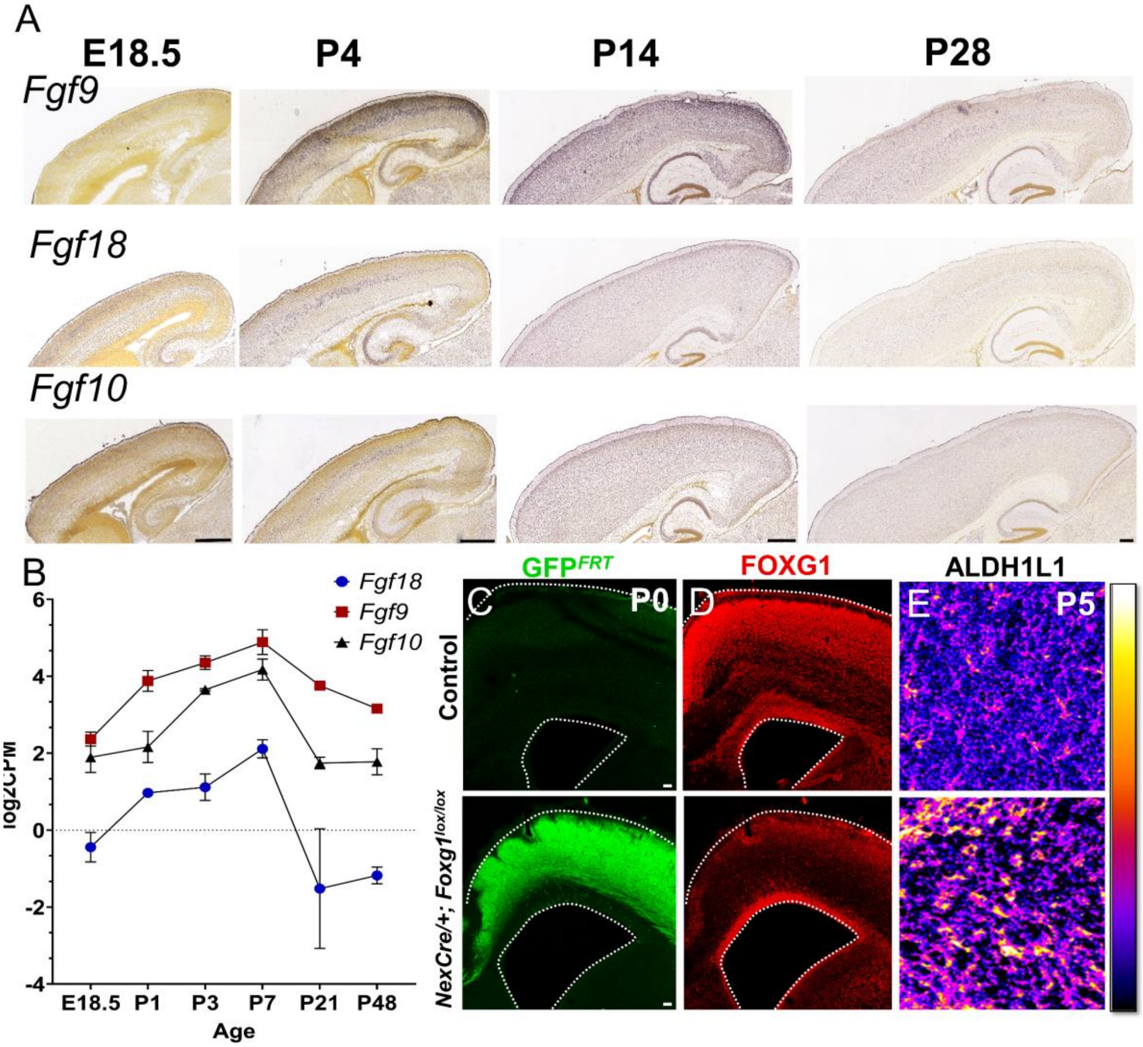


**Supplementary figure S6: Postmitotic neuron-specific role of FOXG1 in regulating FGF signalling.** (A, B) Expression of FGF ligands such as *Fgf9, Fgf10*, and *Fgf18* peaks in the first postnatal week and declines thereafter as seen in Allen Brain Atlas *in situ* hybridization data (A; (53)) and RNA-Seq data from Yuan *et al.*, (52). (C–E) Deletion of *Foxg1* using *NexCre* results in a reduction of FOXG1 protein expression specifically within the cortical plate (C, D) and an apparent increase in ALDH1L1 staining within the cortical plate (E).

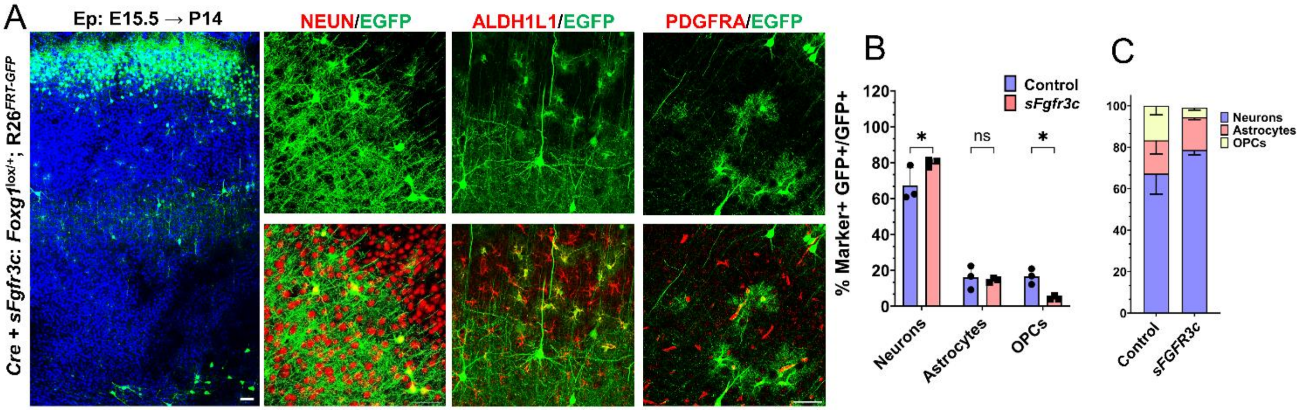


**Supplementary figure S7: *sFgfr3c* overexpression at E15.5 in *$R26^{FRT\text{-}GFP}$* background leads to prolonged neurogenesis.**

(A–C) *sFgfr3c* + *Cre* electroporation at E15.5 in Control (A, *$Foxg1^{lox/+}$*; *$R26^{FRT\text{-}GFP}$*) embryos, followed by analysis at P14 (A). 79.9% of GFP+ cells colocalise with NEUN, compared to 67.3% in Controls. 14.55% of GFP+ cells colocalise with ALDH1L1 compared to 16% in Controls. 3.9% of cells colocalise with PDGFRA compared to 17% in Controls (B, C). n=2000 cells from N=3 biologically independent experiments. S*tatistical analysis: Student's T-Test. Scale bars: 50 µm.*

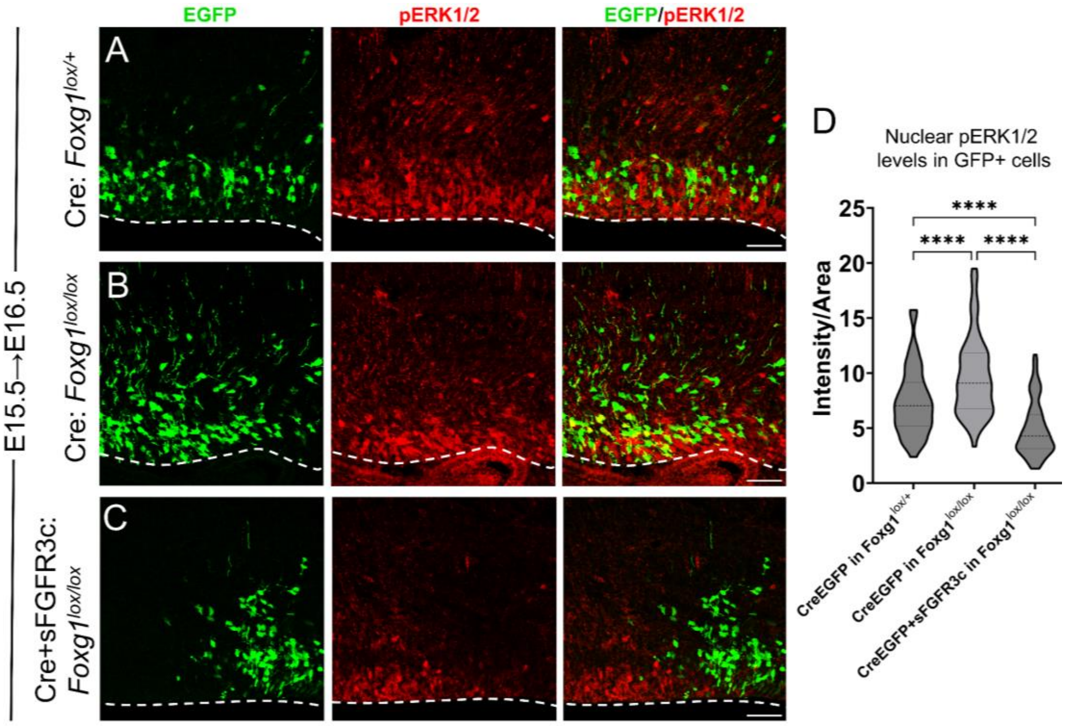


**Supplementary figure S8: FGF Signalling is modulated upon induction of sFGFR3c in E15.5 progenitors.**

(A-D) Quantification of pERK1/2 levels in E15.5 progenitors examined at E16.5, one-day post-electroporation, under different conditions: (A, B) Electroporation of Cre in *Foxg1*$^{lox/lox}$ embryos (B) at E15.5 results in elevated pERK1/2 levels compared to controls (A). This increase is significantly attenuated upon electroporation of Cre+sFGFR3c in *Foxg1*$^{lox/lox}$ progenitors (C). The intensity quantification is represented in D. n=500 cells per condition from N=2 biologically independent experiments. S*tatistical analysis: Two-way ANOVA with Tukey's correction. Scale bars: 50 µm.*

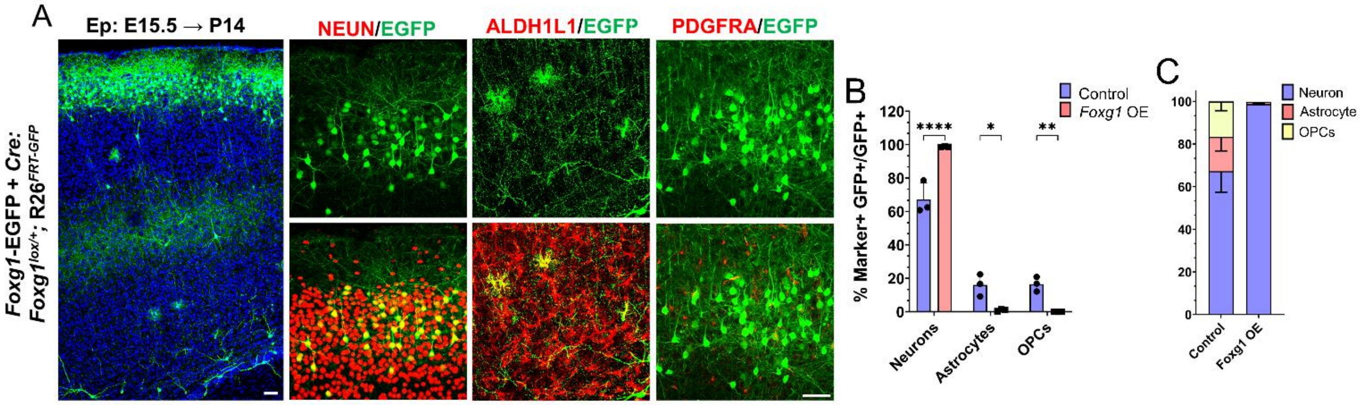


**Supplementary Figure S9: *Foxg1* overexpression at E15.5 in *R26*$^{FRT\text{-}GFP}$ background leads to prolonged neurogenesis.**

(A–C) *Foxg1-Egfp* + *Cre* electroporation at E15.5 in Control (A, *Foxg1*$^{lox/+}$; *R26*$^{FRT\text{-}GFP}$) embryos, followed by analysis at P14 (A). 98% of GFP+ cells colocalise with NEUN, compared to 67.3% in Controls. 2% of GFP+ cells colocalise with ALDH1L1 compared to 16% in Controls. 0% of cells colocalise with PDGFRA compared to 17% in Controls (B, C). n=2000 cells from N=3 biologically independent experiments. *Statistical analysis: Student's T-Test. Scale bars: 50 µm.*

**<u>Supplementary Tables:</u>**

| PRIMARY ANTIBODIES | SPECIES | COMPANY | CATALOGUE NUMBER | DILUTION |
|---|---|---|---|---|
| Biotinylated GFP | Goat | Abcam | ab6658 | 1:200 |
| NEUN | Rabbit | ThermoFisher Scientific | 702022 | 1:200 |
| ALDH1L1 | Rabbit | Abcam | ab87117 | 1:200 |
| OLIG2 | Rabbit | Merck Millipore | AB9610 | 1:200 |
| SOX9 | Rabbit | Abcam | ab185230 | 1:200 |
| KI67 | Rabbit | ThermoFisher Scientific | MA5-14520 | 1:1000 |
| RFP | Mouse | ThermoFisher Scientific | MA5-15257 | 1:200 |
| Phospho-P42/44-MAPK | Rabbit | Cell Signaling Technology | 4370S | 1:200 |
| FGFR3 | Rabbit | Affinity Biosciences | AF0160 | 1:200 |
| NF1A | Rabbit | Abcam | ab228897 | 1:500 |
| PAX6 | Mouse | ThermoFisher Scientific | MA1-109 | 1:500 |
| EOMES | Rat | ThermoFisher Scientific | 14-4875-82 | 1:200 |
| SOX2 | Rabbit | ThermoFisher Scientific | MA1-014 | 1:200 |
| CD140a (PDGFRα) | Rat | BD Biosciences | 558774 | 1:500 |
| FOXG1 | Rabbit | TakaraBio | M227 | 1:200 |

**Supplementary Table S2:**

| SECONDARY ANTIBODIES | SPECIES | COMPANY | CATALOGUE NUMBER | DILUTION |
|---|---|---|---|---|
| Anti-rabbit 568 | Goat | ThermoFisher Scientific | A11011 | 1:200 |
| Streptavidin 488 | Goat | ThermoFisher Scientific | S32354 | 1:200 |
| Anti-rabbit 647 | Donkey | ThermoFisher Scientific | A31573 | 1:200 |
| Anti-mouse 568 | Goat | ThermoFisher Scientific | A11004 | 1:200 |
| Anti-rat 647 | Goat | ThermoFisher Scientific | A21247 | 1:200 |
| Anti-rat 568 | Goat | ThermoFisher Scientific | A11077 | 1:200 |
| Anti-rabbit 488 | Goat | ThermoFisher Scientific | A11034 | 1:200 |

**Dataset S1 (separate file):**

Sheet 1: Differentially expressed genes identified using DESeq2 in the *Foxg1* LOF cells vs. Control at E17.5 related to Figure 3A.

Sheet 2: List intersecting regions between the FOXG1 ChIP-seq and bivalent marks H3K27me3 and H3K4me3.

**Dataset S2 (separate file):**

Sheet 1: Differentially expressed genes identified using DESeq2 in *NexCre/+; Foxg1* LOF vs Control cortical plate cells related to Figure 5E.

Sheet 2: List of differentially expressed secreted molecules subset from the *NexCre/+; Foxg1* LOF vs Control RNA Seq dataset.
